# Genomic characterization of *Vibrio parahaemolyticus* from Pacific white shrimp and rearing water in Malaysia reveals novel sequence types and structural variation in genomic regions containing the *Photorhabdus* insect-related (Pir) toxin-like genes

**DOI:** 10.1101/714626

**Authors:** Chrystine Zou Yi Yan, Christopher M. Austin, Qasim Ayub, Sadequr Rahman, Han Ming Gan

**Author notes:** Corresponding author Name: Han Ming Gan.

## Abstract

The Malaysian and global shrimp aquaculture production has been significantly impacted by acute hepatopancreatic necrosis disease (AHPND) typically caused by *Vibrio parahaemolyticus* harboring the pVA plasmid containing the *pirA*^*Vp*^and *pirB*^*Vp*^genes which code for *Photorhabdus* insect-related (Pir) toxin. The limited genomic resource for *V. parahaemolyticus* strains from Malaysian aquaculture farms precludes an in-depth understanding of their diversity and evolutionary relationships. In this study, we isolated shrimp-associated and environmental (rearing water) *V. parahaemolyticus* from three aquaculture farms located in Northern and Central Malaysia followed by whole-genome sequencing of 40 randomly selected isolates on the Illumina MiSeq. Phylogenomic analysis and multilocus sequence typing (MLST) reveal distinct lineages of *V. parahaemolyticus* that harbor the *pirAB*^*Vp*^genes. The recovery of pVA plasmid backbone devoid of *pirA*^*Vp*^ or *pirAB*^*Vp*^ in some *V. parahaemolyticus* isolates suggests that the toxin genes are prone to deletion. The new insight gained from phylogenomic analysis of Asian *V. parahaemolyticus*, in addition to the observed genomic instability of pVa plasmid, will have implications for improvements in aquaculture practices to diagnose, treat or limit the impacts of this disease.

## Introduction

Acute hepatopancreatic necrosis disease (AHPND), also known as early mortality syndrome, is an emerging shrimp disease that has caused more than 60% reduction in shrimp (*Litopenaeus vannamei*) production worldwide, with global economic losses estimated to be up to USD $1 billion annually (FAO, 2013). Initially reported in China in 2009, this disease continued to spread to other parts of Asia including Vietnam, Thailand, and Malaysia (Zorriehzahra & Banaederakhshan, 2015). To date, AHPND is still prevalent in Malaysia despite a lower frequency of occurrence in recent years (Chu *et al*., 2016).

The causative agent of AHPND was successfully isolated through laboratory assays that fulfill the Koch’s postulates and was subsequently identified as a specific strain of *V. parahaemolyticus* (Tran *et al*., 2013). Comparative genomics of AHPND-causing and avirulent *V. parahaemolyticus* strains identified the presence of a 70-kb pVa plasmid containing the *pirAB*^*Vp*^genes in the virulent strain that code for binary toxins with structural similarity to the *Bacillus* Cry insecticidal toxin-like proteins (Lee *et al*., 2015). This binary toxin induces cell death through pore formation in shrimp cell membranes leading to tissue degradation and digestive organ dysfunction (Lai *et al*., 2015, Lee *et al*., 2015, Han *et al*., 2016). Current genomic sampling suggests that most of AHPND-causing *Vibrio* spp. belong to the species *V. parahaemolyticus* that cluster within the *Vibrio harveyi* clade (Tran *et al*., 2013). However, in recent years, the pVa plasmid has also been isolated from other *Vibrio* species thus indicating the mobility and compatibility of the virulent plasmid among members of the *Vibrio* genus (Kondo *et al*., 2015, Han *et al*., 2016, Dong *et al*., 2017, Xiao *et al*., 2017, Liu *et al*., 2018).

The increasing genomic representation of *Vibrio* sequences in public database enables a more comprehensive analysis and understanding of the evolutionary relationships among different species and strains (Osama *et al*., 2012, Jun *et al*., 2013, Gomez-gil *et al*., 2014, Gomez-Jimenez *et al*., 2014, Kondo *et al*., 2014, Yang *et al*., 2014, Castillo *et al*., 2015, Yang *et al*., 2015). The generation of high-quality draft genomes also allows the extraction of full-length house-keeping genes used for multilocus sequencing typing (MLST), precluding the laborious steps associated with amplification and sequencing of multiple PCR fragments (Osama *et al*., 2012, Joensen *et al*., 2014, Kavousi *et al*., 2016). Successful MLST of the *V. parahaemolyticus* genomes has been recently demonstrated in a comparative genomic analysis of 93 publicly available *V. parahaemolyticus* genomes and 9 newly sequenced Chinese *V. parahaemolyticus* isolates (Fu *et al*., 2017). A recent study identified substantial genomic diversity between outbreak countries and indicated at least 12 independent AHPND-related genomic clones based on whole-genome SNP analysis (Fu *et al*., 2017). Despite the comprehensive genomic sampling, nearly half of the strains analyzed are not associated with AHPND and of the 9 newly sequenced Chinese isolates, only 2 strains were associated with AHPND (Fu *et al*., 2017). Samples size calculations based on Bayesian method of inference recommended that 2 to 20 isolates per sample source should be characterized to obtain maximal information on strain heterogeneity, which is crucial for accurate prediction of outbreak sources and transmission pathways (Döpfer *et al*., 2008). Therefore, it remains unclear the degree to which insufficient sampling may have affected the accuracy of AHPND epidemiological predictions. To date, such a sample size requirement remain difficult to fulfill in genome-based epidemiology of aquaculture-associated pathogens despite the decreasing cost associated with microbial genome sequencing (Land *et al*., 2015, Tang *et al*., 2017).

Genomic resources for bacterial isolates from Malaysia are scarce and were generally presented as individual genome report without a strong focus on comparative genomics (Hong *et al*., 2012, Yap *et al*., 2012, Gan *et al*., 2017). For example, in 2017, only five genomes of Malaysian *V. parahaemolyticus* associated with shrimp aquaculture environment were reported (Foo *et al*., 2017) followed closely by only two additional pVa-harbouring *V. parahaemolyticus* genomes a year later, each presented as a single genome report (Devadas *et al*., 2018, Devadas *et al*., 2018). Such a small dataset (n < 10) limits the inference of the genomic diversity and evolutionary relationships of *V. parahaemolyticus* in Malaysia. In this study, we attempt to overcome this shortcoming by sequencing an additional 40 *V. parahaemolyticus* isolates from deceased shrimps and rearing water collected from three Malaysian shrimp aquaculture farms. From the *de-novo* assembled genomes, we characterized their genetic diversity based on *in-silico* MLST and determined their evolutionary relationships with other Asian *V. parahaemolyticus* isolates. The genetic and structural analysis of the pVa plasmids was also investigated, providing new insights into their prevalence and diversity among Asian *V. parahaemolyticus* strains.

## Materials and methods

### Field Sampling

Sampling was performed in three Malaysian shrimp aquaculture farms with suspected AHPND outbreaks. Two of these were located in Negeri Sembilan, Central Peninsular Malaysia and one in Terengganu, Northern Peninsular Malaysia (Figure 1). In all three aquaculture farms, pond rearing water was collected for strain isolation. Two deceased shrimps exhibiting the symptoms of AHPND such as empty stomach, pale hepatopancreas, and empty midgut were also provided by the farm manager of the Terengganu (Northern Malaysia) aquaculture farm (Table 1). At the request of the aquaculture farm managers, the exact sampling locations were not disclosed in this study to protect the identity of the farms.

**Figure 1:**
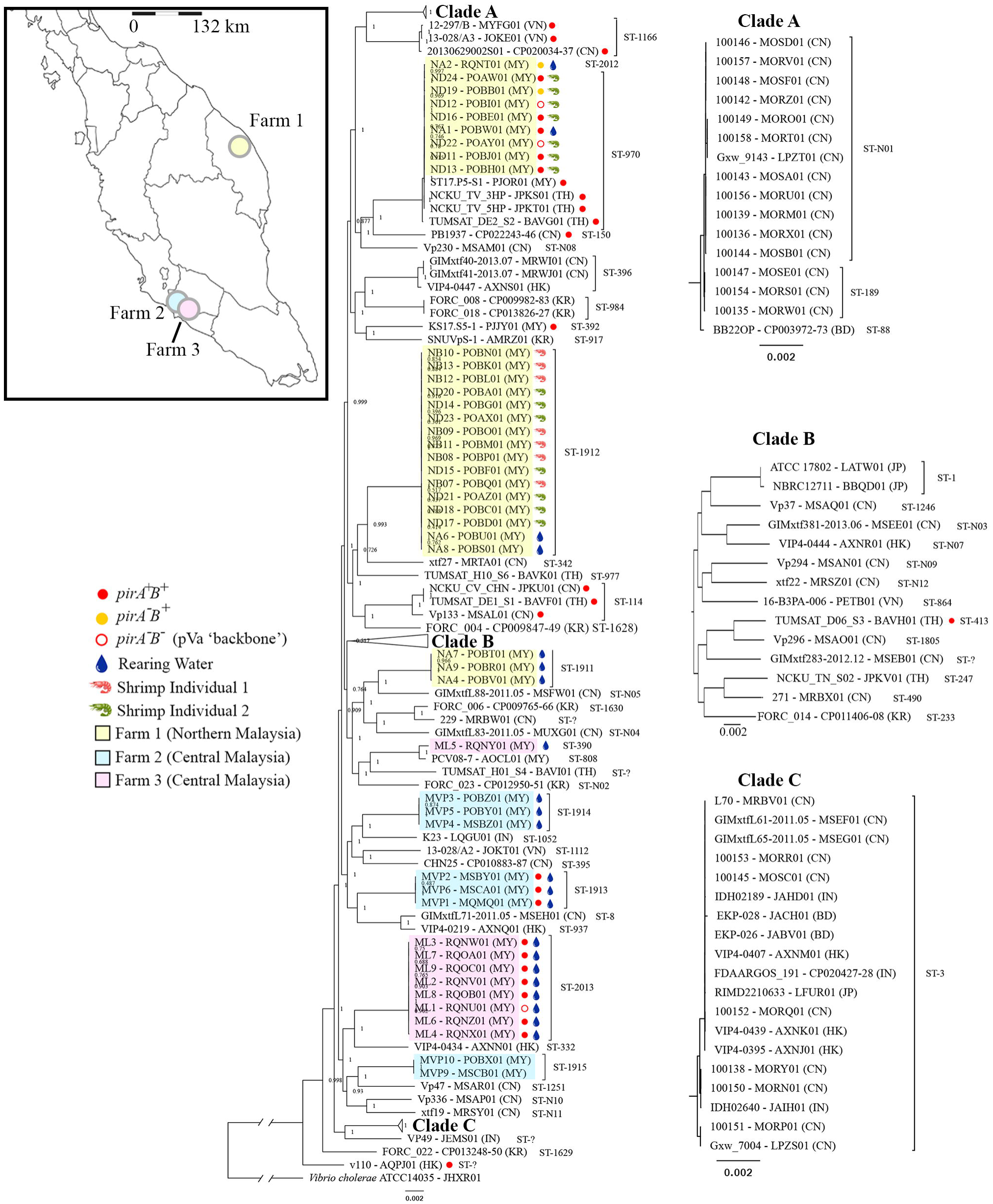
Maximum likelihood tree based on core genome alignment depicting the evolutionary relationships of 135 Asian *Vibrio parahaemolyticus* isolates. Approximate sampling location was indicated in the peninsular Malaysia map located at the top left corner. Sequence types are labeled accordingly with ST followed by their designated profile number based on PubMLST database (https://pubmlst.org/vparahaemolyticus). The tree was rooted with *V. cholera* as the outgroup and for clarity, the root has been shortened by approximately 80% of the actual length shown. Branch lengths and node values indicate the number of substitutions per site and SH-like support values, respectively. Two-letter codes in brackets next to the tip label indicate sample origin. MY, Malaysia; HK, Hong Kong, CN, China; KR, Korea; IN, India; VN, Vietnam; TH, Thailand.

### Strain isolation and DNA extraction

Enrichment of *Vibrio* spp. from homogenized shrimp hepatopancreas and pond water was performed on thiosulfate-citrate-bile salts-sucrose (TCBS) agar (Oxoid Ltd, UK). We chose the hepatopancreas as the isolation source for shrimp as it is usually targeted by AHPND *V. parahaemolyticus* as previously determined based on PCR and histopathology (Khimmakthong & Sukkarun, 2017). After incubation overnight at 30°C, blue-green colonies were isolated and sub-cultured on a fresh batch of TCBS agar followed by similar incubation setting, to verify their identity as *V. parahaemolyticus*. The confirmed blue-green colonies were sub-cultured onto nutrient agar (Merck, Kenilworth, NJ, USA) supplemented with 3% (w/v) NaCl. These culture plates were incubated overnight at 30°C and DNA from pure single colonies were extracted using a modified salting-out method (Sokolov, 2000).

### Whole-genome sequencing

DNA from each isolate was normalized to 0.2 ng/uL based on Qubit2 measurement (Thermo Fisher Scientific, Waltham, MA, USA) and used to prepare NexteraXT sequencing library (Illumina, San Diego, CA, USA) according to the manufacturer’s instructions. Each constructed library contains a unique combination of Nextera dual-index barcodes that allow pooling and sequencing on the same MiSeq flowcell. Equal molarity of each barcoded library was pooled followed by preparation for sequencing on the Illumina MiSeq sequencer located at Monash University Malaysia (2×250 bp paired-end read setting).

### *De novo* assembly and *in-silico* taxonomic validation

Raw MiSeq read files were filtered for adapter sequences and low-quality base calls using Trimmomatic version 0.36 (Bolger *et al*., 2014) with the parameters PE, ILLUMINACLIP:Trimmomatic/adapters/NexteraPE.fa:2:30:10, LEADING:3, TRAILING:3, SLIDINGWINDOW:4:20, MINLEN:20. The trimmed reads were then assembled into contigs using SPAdes version 3.10.0 (Bankevich *et al*., 2012) with the default parameters. The assembled contigs were subsequently scaffolded and gap-closed using SSPACE version 3.0 (Boetzer *et al*., 2011) and GapFiller v1.10 (Boetzer *et al*., 2012) respectively. QUAST-version 4.4 (Gurevich *et al*., 2013) was used to assess the assembly using *V. parahaemolyticus* ATCC 17802 ^T^ as a reference genome. To confirm species identity, JSpecies version 1.2.1 (Richter *et al*., 2015) was used to calculate the pairwise average nucleotide identity (ANI) of the assembled genomes with *V. parahaemolyticus* ATCC 17802^T^ as the positive control and *Vibrio cholera* ATCC 14035^T^ as the negative control.

### Genome annotation and multi-locus sequence typing

Identification of pVa plasmid, *pirA*Vp and *pirB*Vp genes was performed using local BLASTN version 2.7.1+. Gene prediction and annotation was performed in NCBI and *in-silico* MLST was based on seven housekeeping genes in the MLST database (https://pubmlst.org/vparahaemolyticus/: *dnaE, dtdS, gyrB, pyrC, pntA, recA*, and *tnaA*) using three approaches e.g. local BLASTN version 2.7.1+, SRST-2 version 0.2.0, and MLST 1.8 (Camacho *et al*., 2009, Larsen *et al*., 2012, Inouye *et al*., 2014).

### Whole-genome phylogeny

Publicly available *V. parahaemolyticus* genomes were downloaded from NCBI Assembly in October 2018. Genomes with poor assembly metrics (> 400 contigs), inconsistent sample origin or erroneous taxonomic assignment were removed (Supplementary Table 1). For the identification of pan-genome, core-genome and accessory genome from the filtered and forty newly assembled *V. parahaemolyticus* genomes (Table 1), we used Roary version 3.12.0 (Page *et al*., 2015) and decreased the minimum BLASTP percentage identity from 95% to 90%. In addition, the *V. cholerae* ATCC 14035^T^ genome was also included in the Roary analysis as the outgroup. A maximum-likelihood tree was then constructed based on the alignment of the core-genomes using FastTree version 2.1.8 (Price *et al*., 2010) with generalized time-reversible (GTR) model. Visualization of the reconstructed maximum likelihood tree used FigTree v1.4.3.

### Plasmid and gene neighborhood visualization

The presence of pVA among the *V. parahaemolyticus* genome assemblies was assessed based on BLASTN search (e-value of 1e-5) against the reference pVa from *V. parahaemolyticus* 13-028/A3 (GenBank Accession Number: NC_025152). Nucleotide alignment and visualization of the selected pVa-harbouring *V. parahaemolyticus* genomes against reference pVa used BRIG version 0.95 (Alikhan *et al*., 2011). Visualization of the gene neighborhood surrounding the *pirB*^Vp^ among *pirA*^Vp-^ *pirB*^Vp+^ isolates used EasyFig version 2.1 with BLASTN search (e-value of 0.001, minimum length coverage of 100 bp) (Sullivan *et al*., 2011).

## Results

### Genomic characterization and *in-silico* taxonomic validation of sequenced strains

We sequenced and assembled forty whole-genomes of *V. parahaemolyticus* isolates from three farms in Peninsular Malaysia using the Illumina MiSeq platform. The assembled whole-genomes of these *V. parahaemolyticus* strains showed an average cumulative length of 5 Mb and 45% GC content, consistent with the assembly statistics of most published genomes of *V. parahaemolyticus* in the NCBI database (Supplementary Table 1) (Jun *et al*., 2013, Gomez-gil *et al*., 2014, Gomez-Jimenez *et al*., 2014, Kondo *et al*., 2014, Yang *et al*., 2014, Castillo *et al*., 2015, Kondo *et al*., 2015, Yang *et al*., 2015). All newly sequenced isolates exhibited more than 95% pairwise average nucleotide identity (ANI) to *V. parahaemolyticus* ATCC 17802 (Supplementary Figure 1>).

### High genomic diversity of *Vibrio parahaemolyticus* associated with Malaysian aquaculture farms

Phylogenomic clustering of isolates corresponds broadly with their MLST profiles instead of their country of isolation. The 40 Malaysian *V. parahaemolyticus* isolates sequenced in this study as well as 5 from our previous work (Foo et al., 2017) formed 8 distinct genomic clusters of which 6 consist entirely of rearing water isolates (Figure 1). Based on the current genomic sampling, with the exception for the genomic cluster corresponding to ST-970, the remaining 7 genomic clusters are unique to Malaysian aquaculture farm isolates with each cluster consisting of only isolates that are exclusive to a specific farm (Figure 1).

### New sequence types of *Vibrio parahaemolyticus* revealed by *in-silico* multilocus sequence typing

MLST profiling indicates that that the *dtdS* gene coding for threonine 3-dehydrogenase is the most variable among the 7 housekeeping genes of currently sequenced Asian *V. parahaemolyticus* with 45 alleles identified (Supplementary Table 1). Conversely, the *pntA* gene coding for NAD(P) transhydrogenase subunit alpha subunit has the least number of observed alleles (n=28, 5 are found in the isolates of this study) (Supplementary Table 1). A total of 33 known and 18 new sequence types (STs) were identified from the currently sequenced Asian *V. parahaemolyticus* strains of which 7 (ST-1911 to 1915, ST-2012 and ST-2013) were exclusive to the *V. parahaemolyticus* isolates in this study (Figure 1). All currently sampled *V. parahaemolyticus* isolates that were assigned to ST-390, ST-1911, ST-1912, ST-1914, ST-1915 contain no detectable *pirAB*^*Vp*^ and pVa plasmid backbone in their genome assemblies. Among the pVa-harbouring Malaysian isolates, a majority of them were assigned to ST-970 (n=9) followed by ST2013 (n=8), ST-1913 (n=3), ST-2012 (n=1) and ST-392 (n=1) which together formed four major genomic clusters in the phylogenomic tree (Figure 1).

### Three major structural variants of the pVA plasmid among Asian *V. parahaemolyticus*

Of the 52 sequence types of *V. parahaemolyticus* identified in this current genomic sampling, *pirAB*^*Vp*^ could only be detected among isolates from 8 validated (ST-2013, ST-114, ST-150, ST-392, ST-413, ST-970, ST-1166, and ST-1913) and 1 undetermined sequence types (Figure 1). Strain v110 with the undetermined sequence type has an unassigned *pntA* allele with the closest match to *pntA48* (Supplementary Table 1). Whole-genome alignment of pVA-harbouring Asian *V. parahaemolyticus* isolates against the reference pVA plasmid showed near-complete sequence conservation of the plasmid except for the *pir*^*Vp*^ gene region that exhibits three major structural variants e.g. *pirA*^*Vp*+^*B*^*Vp*+^, *pirA*^*Vp*-^*B*^*Vp*+^ and *pirA*^*Vp*-^*B*^*Vp*-^ (Figure 2 and Supplementary Figure 2). A closer inspection of the *pirB* gene neighborhood in both *pirA*^*-*^*B*^*+*^ NA2 and ND19 mutants revealed a complete deletion of the *pirA*^*Vp*^ gene and a slight truncation at the 5’ end of *pirB*^*Vp*^. Immediately upstream of the 5’ truncated *pirB*^*Vp*^ gene is a small fragment of a transposase gene showing significant nucleotide homology to that of the reference pVa plasmid (Figure 3). The observed gene organization in NA2 and ND19 contrasted with that of *V. parahaemolyticus* mutant isolate XN87 that appears to have a transposase inserted into the *pirA*^*Vp*^ gene (Figure 3).

**Figure 2:**
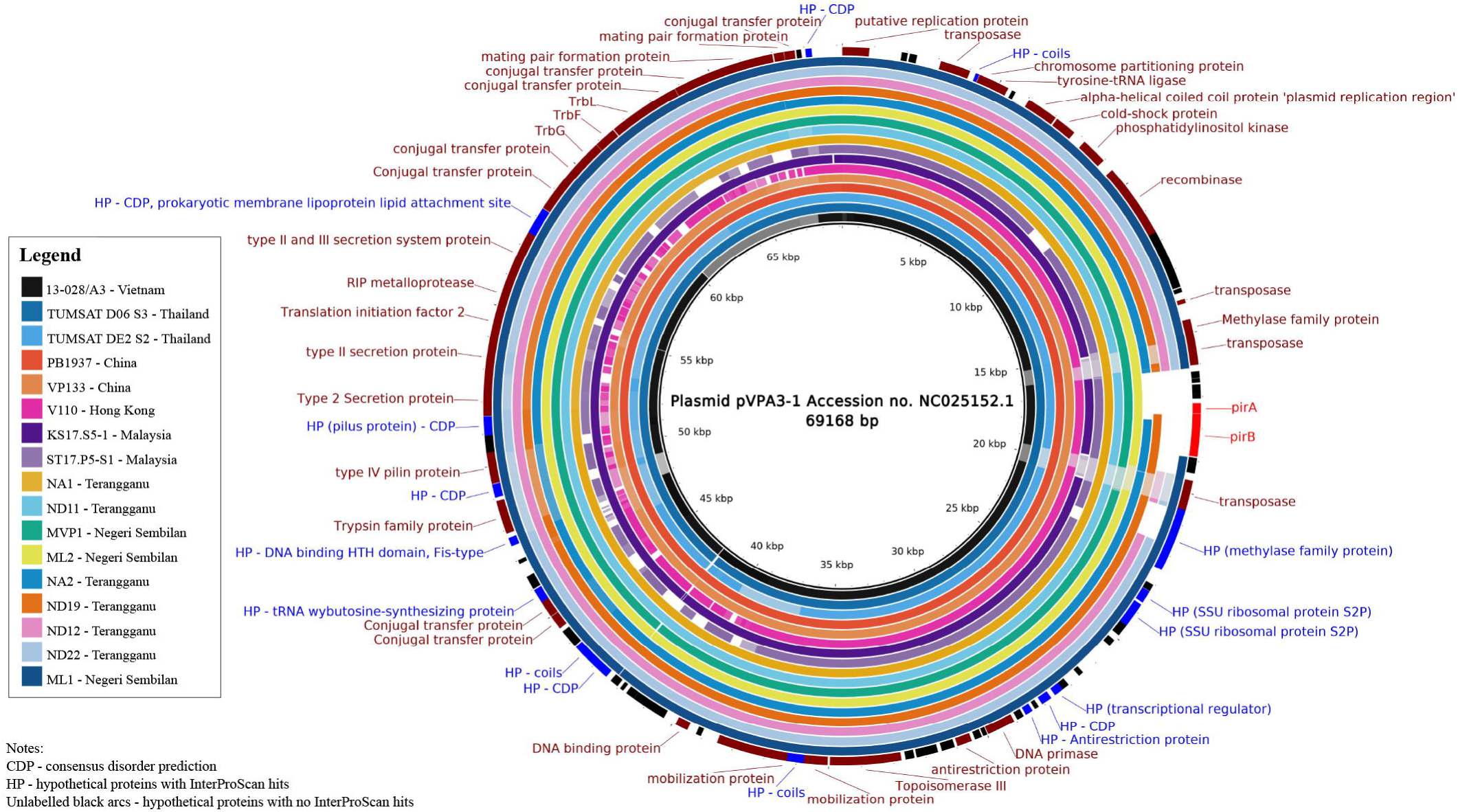
BRIG visualization of *V. parahaemolyticus* pVa plasmid. Using BLASTN analyses, pVa plasmid from clade-representative strains of *V. parahaemolyticus* throughout the Asian region were aligned against plasmid pVPA3-1 and visualized using BRIG. Black arcs indicate hypothetical proteins with no InterProScan hits. The innermost circle represents the reference sequence of plasmid pVPA3-1 harbored by *V. parahaemolyticus* strain 13-028/A3. Outer rings illustrate shared identity with pVa plasmids found in some samples of this study and NCBI database. The varying saturation of the ring colors is based on 100%, 90% and 70% identity. Clear white gaps indicate no sequence similarity in that area. Full visualization of all *V. parahaemolyticus* strains harboring pVa plasmid can be found in Supplementary Figure 2.

**Figure 3:**
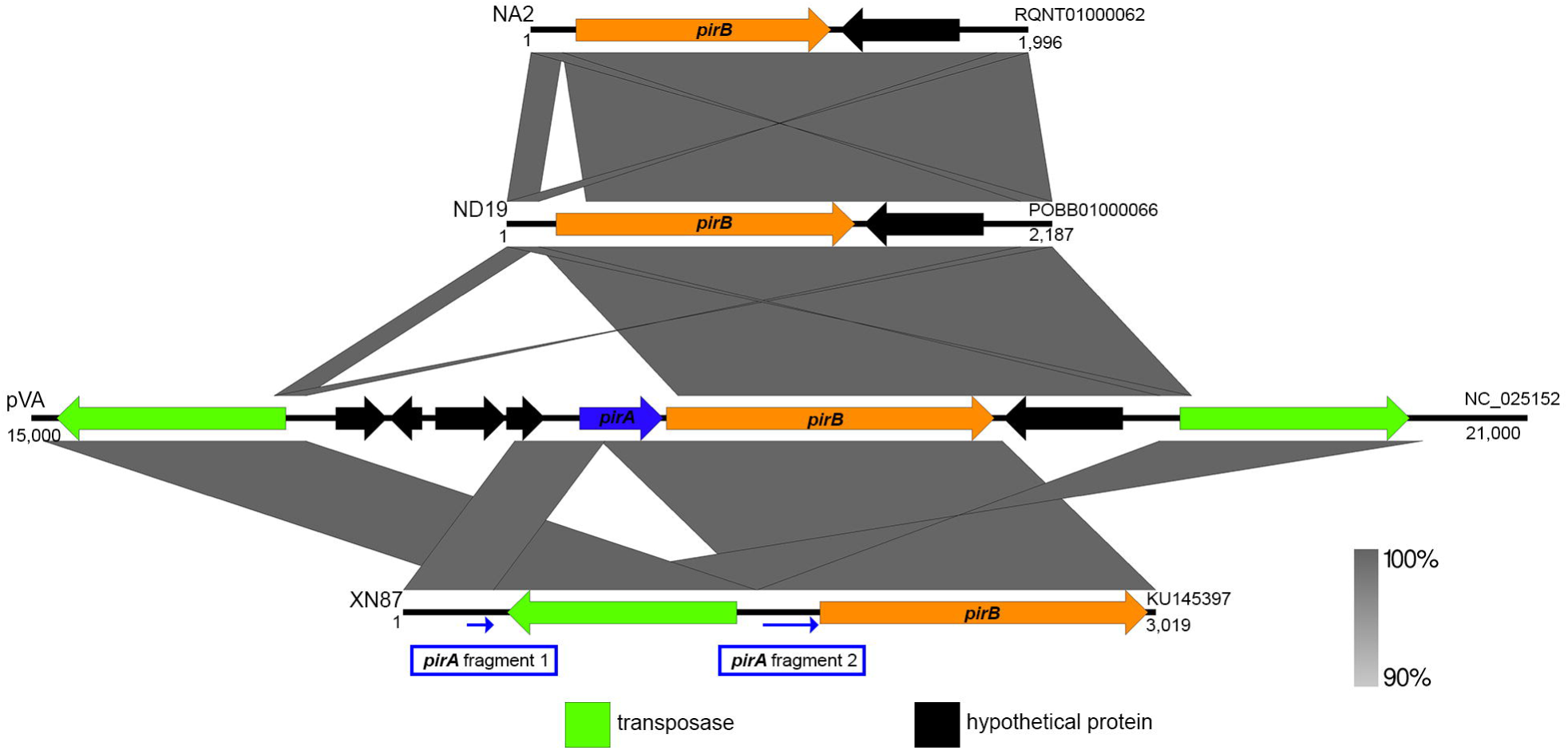
Gene organization of *pirB*^*Vp*^-containing contigs among selected *V. parahaemolyticus* isolates. Vertical blocks between sequences indicate regions of homology shaded according to BLASTn. Left and right labels on each sequence are isolate names and GenBank accession numbers, respectively. Blue arrows below the XN87 sequence indicate the location of transposase-inserted pirA genes (*pirA* fragment 1 and *pirA* fragment2) that were not annotated in the original GenBank file. The direction of arrow indicates gene transcription orientation.

## Discussion

We report the whole-genome sequencing of 40 *V. parahaemolyticus* isolated from Pacific white shrimp hepatopancreas and aquaculture rearing water in Malaysia. The phylogenomic clustering indicates at least four distinct origins of *pirAB*^*Vp*^-harbouring *V. parahaemolyticus* in Malaysia pending future sampling and genome sequencing. Multiple origins of *pirAB*^*Vp*^+ *V. parahaemolyticus* were also observed in a previous MLST study in Thailand indicating high genetic diversity among AHPND *V. parahaemolyticus* isolates (Zorriehzahra & Banaederakhshan, 2015). Furthermore, we have substantially expanded the genomic representation within the ST-970 clade, which now consists of *pirA*^*Vp*+^*B*^*Vp*+^, *pirA*^*Vp*-^*B*^*Vp*+^ and *pirA*^*Vp*-^*B*^*Vp*-^ isolates.

In this study, despite the detection of several STs among the *V. parahaemolyticus* isolates with some even sharing the same isolation source, the pVa plasmid could only be detected in isolates from a certain sequence type, suggesting that the distribution of virulence factor for AHPND among culturable *V. parahaemolyticus* maybe ST-specific. Given that a majority of the isolates sequenced in this study were collected from the rearing water, it is also possible that the pVa plasmids were lost due to the lack of selection pressure or had not been acquired and propagated among certain sequence types. A successful AHPND strain must first have genomic makeup that permits replication and maintenance of the pVa plasmid (Petersen, 2011). Secondly, it must be able to colonize and persist in its shrimp host as well as exhibit timely expression of the *pir* genes during pathogenesis (Gode □Potratz *et al*., 2011, Liu & Chen, 2015, Soonthornchai *et al*., 2015, Li *et al*., 2017, Ng *et al*., 2018). Further, given that AHPND leads to nearly 100% host mortality, it is also essential that the genomic make-up of an AHPND strain supports persistence outside its animal host post-death e.g. pond rearing water, which will have a diversity of microbial, chemical and physical compositions (Chen *et al*., 2017, Cornejo-Granados *et al*., 2017, Dittmann *et al*., 2017, Zoqratt *et al*., 2018). That said, the presence of pVa-harbouring Asian *V. parahaemolyticus* in multiple distantly-related clades suggests that many existing *V. parahaemolyticus* may already have the genomic potential to transform into an AHPND strain after the acquisition of the virulent pVa plasmid presumably through conjugal transfer.

The expression of both PirA^*Vp*^ and PirB^*Vp*^ toxins are required to cause the onset of AHPND (Hirono *et al*., 2016), but it has been hypothesized that PirA^*Vp*^ is not expressed until later in the infection stage (Lai *et al*., 2015, Lee *et al*., 2015, Sirikharin *et al*., 2015, Han *et al*., 2016). Interestingly, despite the inability of a *pirA* mutant *V. parahaemolyticus* XN87 to produce the PirA^*Vp*^ and PirB^*Vp*^ toxins or to cause AHPND lesions, it still causes 50% mortality in shrimps presumably from the expression of other putative virulence genes located in the pVA plasmid (Phiwsaiya *et al*., 2017). Instead of a transposon-inserted *pirA*^*Vp*^, we observed a complete *pirA*^*Vp*^ deletion in NA2 and ND19 isolated from the aquaculture farm in Northern Malaysia. Despite the notable structural dissimilarity observed in *pir* region between XN87 and NA2/ND19, the close proximity of the transposase to *pirB*^*Vp*^ in NA2 and ND19 is likely going to disrupt the expression of *pirB*^*Vp*^ due to a polar mutation effect as reported in XN87 and various transposon mutants (Harayama *et al*., 1983, Richardson *et al*., 1990, Gan *et al*., 2011, Gan *et al*., 2012, Phiwsaiya *et al*., 2017).

The *pirAB*^*Vp*^ genes are usually flanked by 2 identical transposons with 18 bp of terminal inverted repeats with significant homology to the IS903 in *Escherichia coli* (Xiao *et al*., 2017). Minor structural variations observed within the pVa region containing the *pirAB*^*Vp*^ genes suggest that they are prone to deletion or transposition. This is supported by previous studies reporting the natural loss of *pirAB*^*Vp*^ genes in the virulent strains of *V. parahaemolyticus* (strain E1) and *V. owensii* after laboratory sub-culturing with an estimated 22% loss rate of *pirAB*^*Vp*^ in *V. owensii* (Tinwongger *et al*., 2016, Liu *et al*., 2018). Given that all *V. parahaemolyticus* isolates in this study have gone through at least two sub-culturing to ensure culture purity prior to whole-genome sequencing, it is possible that the *pir*^*Vp*^ gene(s) were lost from the initially *pir+* isolates in the laboratory rather than in their natural environment, due to the lack of selection pressure (Smith & Bidochka, 1998). Future study investigating the stability of *pirA*^*Vp*^ and *pirB*^*Vp*^ gene via PCR among our isolates under different growth conditions will be required. In addition, whole metagenome sequencing will be instructive to more accurately estimate the fraction of naturally occurring pVa plasmid with *pirA*^*Vp*^ and/or *pirB*^*Vp*^ deletion in the shrimp pond water and affected hepatopancreas (Jitwasinkul *et al*., 2016, Krawczyk *et al*., 2018).

## Conclusions

We report the whole-genome sequencing on 40 *V. parahaemolyticus* isolates from deceased shrimps and rearing water collected from three Malaysian shrimp aquaculture farms. The number of new sequence types, MLST alleles and lineages of *V. parahaemolyticus* identified from only three sampling sites in this study underscores the overlooked genomic diversity of culturable *V. parahaemolyticus* associated with shrimp aquaculture environment. In addition to expanding the genomic resources of *pirAB*^*Vp*^-bearing *V. parahaemolyticus*, especially for sequence types without genomic representation, it is recommended that future studies also include sequencing of avirulent *V. parahaemolyticus* isolates from APNHD-affected shrimp aquaculture environment to gain a better understanding of *V. parahaemolyticus* genomic diversity that may provide new insights into future pathogen management in shrimp aquaculture.

## Acknowledgments

This study is supported by the Ministry of Higher Education under the grant FRGS/1/2016/STG05/MUSM/03/1. We would like to thank Lab-Ind Resource Sdn Bhd for their kind assistance in the sampling process.

## Figure Legends

**Supplementary Figure 1**: JSpecies results based on ANIm calculation with *V. parahaemolyticus* ATCC 17802 as positive control and *V. cholerae* ATCC 14035 as the negative control.

**Supplementary Figure 2**: Compilation of *V. parahaemolyticus* pVa plasmid. Using blastn analyses, *pir* plasmid from strains of *V. parahaemolyticus* throughout the Asian region were aligned against pVPA3-1 and visualized using BRIG. Black arcs indicate hypothetical proteins with no InterProScan hits. The innermost circle represents the reference sequence of plasmid pVPA3-1 identified in *V. parahaemolyticus* strain 13-028/A3. Outer rings illustrate shared identity with Pir plasmids found in some samples of this study and NCBI database.

**Supplementary Table 1**: Summary of strains used in this study with accession number, origin, genome statistics, MLST profiles, and presence of *pirA*^*Vp*^ and *pirB*^*Vp*^.

