## Supplementary figures and images for "Genomic characterization of *Vibrio parahaemolyticus* from Pacific white shrimp and rearing water in Malaysia reveals novel sequence types and structural variation in genomic regions containing the *Photorhabdus* insect-related (Pir) toxin-like genes"

### Supplementary Figure 1

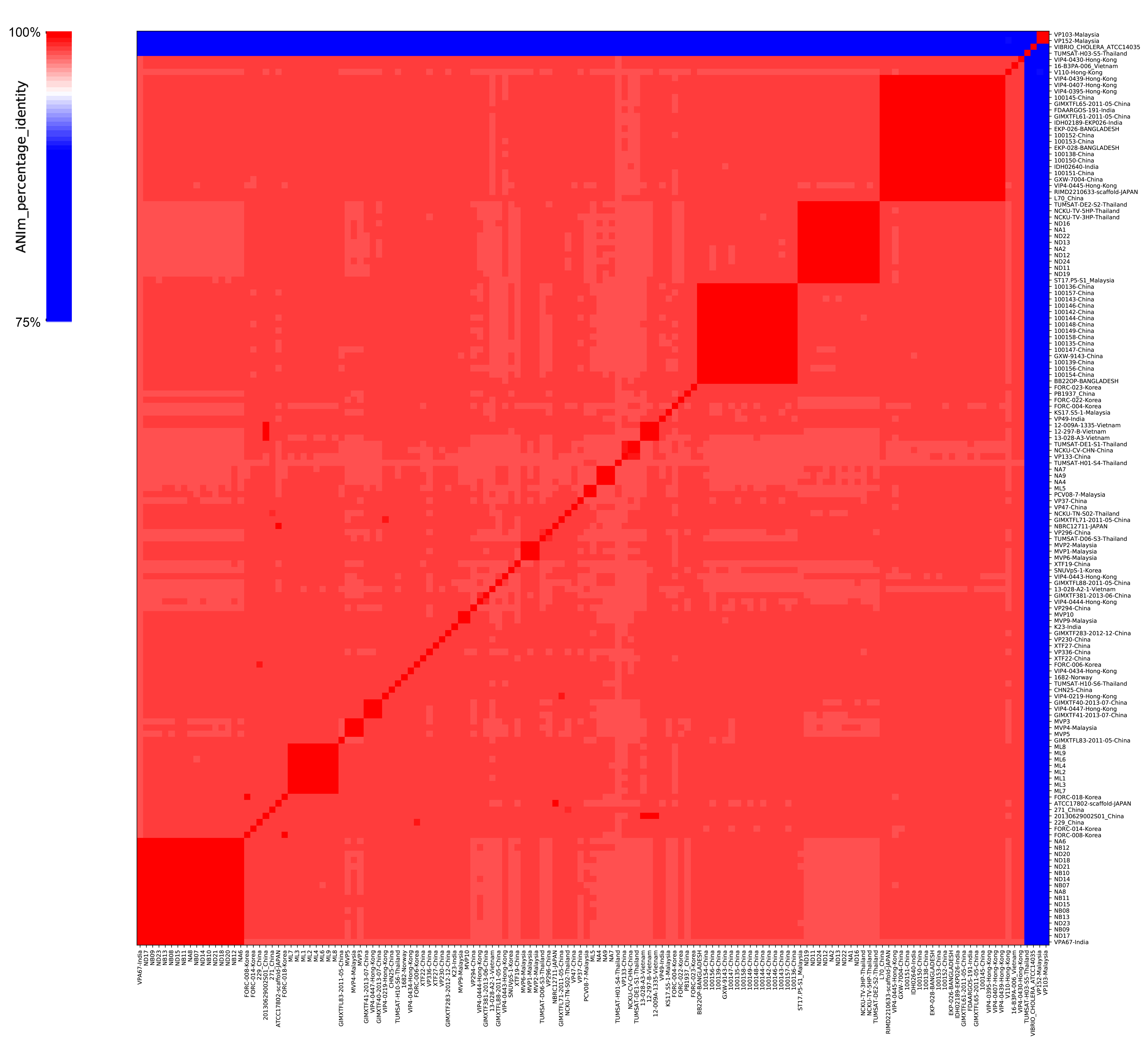

### Supplementary Figure 2

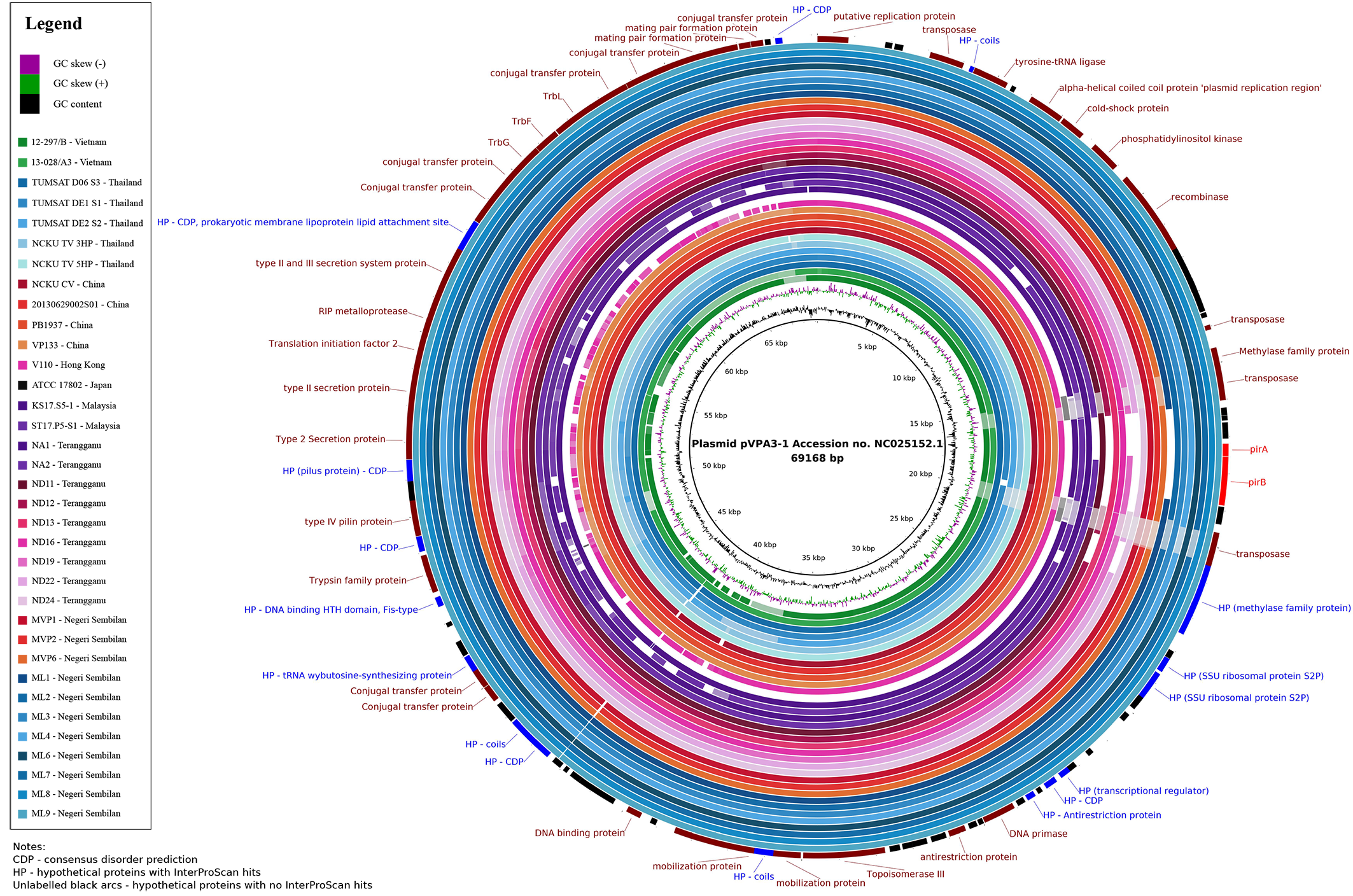
